# Selective attention reduces responses to relevant sounds in human auditory cortex

**DOI:** 10.1101/2022.09.12.507664

**Authors:** Agustin Lage-Castellanos, Federico De Martino, Geoffrey M. Ghose, Omer Faruk Gulban, Michelle Moerel

## Abstract

Selective attention enables the preferential processing of relevant stimulus aspects. Invasive animal studies have shown that attending a sound feature rapidly modifies neuronal tuning throughout the auditory cortex. Human neuroimaging studies have reported enhanced auditory cortical responses with selective attention. To date, it remains unclear how the results obtained with functional magnetic resonance imaging (fMRI) in humans relate to the electrophysiological findings in animal models. Here we aim to close the gap between animal and human research by combining a selective attention task similar in design to those used in animal electrophysiology with high spatial resolution ultra-high field fMRI at 7 Tesla. Specifically, human participants perform a detection task, while the probability of target occurrence varies with sound frequency. Contrary to previous fMRI studies, we show that selective attention reduces responses to the attended frequencies in those neuronal populations preferring the attended frequency. Through population receptive field (pRF) mapping, we furthermore show that these response reductions are at least partially driven by frequency-induced pRF narrowing. The difference between our results to those of previous fMRI studies supports the notion that the influence of selective attention on auditory cortex is diverse and may depend on context, task, and auditory processing stage.

## Introduction

Selective attention highlights currently relevant information through the preferential processing of stimulus features (Maunsell and Treue 2006; Carrasco 2011). Selective attention results in a behavioral advantage (e.g., increased detection performance, faster reaction times) for the attended feature, thereby allowing us to efficiently handle the potentially overwhelming amount of sensory information in our environment.

The neural correlates of selective attention have been extensively investigated throughout stimulus modalities (e.g., in the visual system (Martinez-Trujillo and Treue 2004); somatosensory system (Schweisfurth et al. 2014); and auditory system (Fritz et al. 2003; Lee and Middlebrooks 2011)). In the auditory system, selective attention has also been shown to substantially influence neuronal processing. However, while across-species evidence from the visual system almost uniformly supports a gain model of selective attention (Tootell et al. 1998; Saenz et al. 2002; Hopf et al. 2004; Warren et al. 2014), results from the auditory cortex are diverse. Invasive electrophysiological studies in animals have shown that selectively attending a specific sound feature (e.g., frequency, or spatial location) induces rapid changes in the preference and selectivity of auditory neurons (Fritz et al. 2003, 2007; Fritz 2005; Lee and Middlebrooks 2011; Lakatos et al. 2013; O’Connell et al. 2014). These changes are stronger in secondary and tertiary regions (i.e., belt and parabelt, respectively) than in the primary auditory cortex (Atiani et al. 2014), and are strongest in neurons whose preference matches the attended feature (Atiani et al. 2014). However, these changes are highly task dependent. For example, decreased, instead of increased responsiveness was observed when animals were rewarded for target detection instead of punished for missing a target (David et al. 2012). These results support a matched filter model of selective attention in which auditory neurons change tuning to optimally process attended features (Fritz et al. 2007), but for which the exact changes are dependent on task structure, difficulty, and associated reward (Atiani et al. 2009; David et al. 2012).

By contrast, human functional magnetic resonance imaging (fMRI) studies of auditory selective attention consistently support a gain model in which overall responsiveness (Paltoglou et al. 2009; Da Costa et al. 2013a; Riecke et al. 2017), but not tuning (Dick et al. 2017; Riecke et al. 2018), changes. This discrepancy with results from animal electrophysiology could in part be due to the limited spatial resolution of fMRI. While changes in voxel tuning can be reliably assessed with fMRI (Kay et al. 2015), the voxel response could obscure changes at the neuronal level (Sadil et al. 2021). Conflicting findings between human fMRI and non-human invasive studies may also have been caused by their differences in experimental design. While target and distractor sounds were presented sequentially in animal electrophysiology studies, in human fMRI studies target and distractor sounds were always presented simultaneously, i.e., in an auditory scene (note that while Da Costa et al. (2013b) included a “single stream” condition, they did not evaluated the effect of attention within this condition). Finally, the fMRI studies evaluated frequency tuning with limited spectral resolution (Dick et al. 2017; Riecke et al. 2018) and therefore might be less sensitive to subtle changes. As a result, to date it is still largely unclear how the results obtained with fMRI in humans relate to the electrophysiological findings in animal models.

Here we employed a selective attention task where participants alternatively directed their attention to low or high frequencies. By presenting target and distractor sounds sequentially and using ultra-high field fMRI, we aimed to close the gap between animal and human research. We measured natural sounds responses to estimate population receptive field changes with high spectral resolution. Contrary to previous fMRI studies we observed a response reduction to attended sounds, which resulted at least in part from attention-induced changes in voxel tuning. Our results, and their difference with results from previous human fMRI investigations, thereby support the notion arising from animal studies that, depending on context, task, and auditory processing stage, diverse mechanisms underlie auditory selective attention.

## Materials and methods

### Ethics

The experimental procedures were approved by the Ethics Review Committee of the Faculty of Psychology and Neuroscience at Maastricht University (#167_09_05_2016_S2). The experiment was performed in accordance with the approved guidelines and the Declaration of Helsinki. Informed consent was obtained from each participant before starting the measurements. Participants received course credit or gift vouchers for their participation.

### Participants

Eight healthy volunteers, without history of hearing disorder or neurological disease, participated in this study (mean age [SD] = 28.1 [3.2]; four males and four females). A pure-tone audiogram (with a 25 dB hearing level threshold) was conducted to ensure the participants did not have hearing loss.

### Experimental design and statistical analysis

#### Noise burst detection task

Selective attention was manipulated through a noise-burst detection task on artificial ripple sounds. Specifically, participants were presented with ripples centered at a low or high sound frequency (two center frequencies [300 Hz; 4 kHz] × two temporal modulation rates [3 Hz; 10 Hz]; ripple bandwidth = 1 octave; modulation depth = 0.6; duration = 1 s, with 50 ms onset and offset linearly ramped) and were instructed to press a button when they detected a short white noise burst (duration = 112 ms, with 6 ms onset and offset linearly ramped) in the ripple sounds. The noise burst occurred 500 - 800 ms after ripple onset, and always started at a peak intensity of the ripple envelope. The intensity of the noise burst was individually calibrated to ensure equal detection difficulty across ripple center frequencies and equal performance across participants (see below for details). Participants received feedback about their performance by a change in the color of the fixation cross (from black to either green [correct] or red [incorrect]) upon button press. By manipulating the probability of noise burst occurrence, it was made advantageous to either attend low frequency (300 Hz) or high frequency (4 kHz) ripple sounds. Specifically, in the “Attend Low” condition, 70% of the ripples with a center frequency of 300 Hz contained a noise burst, while the noise burst was present in only 30% of the ripples with a center frequency of 4 kHz. In the “Attend High” condition, the percentage of ripples containing a noise burst was reversed (30% of the 300 Hz ripples, and 70% of the 4 kHz ripples). The attentional condition was kept equal throughout a run of several minutes (see below for details), and runs were presented in a randomized order. At the start of each run, participants were informed regarding the sound frequency for which noise bursts were most common. All ripple sounds, with and without noise burst, were generated with Matlab (The MathWorks, Inc.) and presented to the participants using the Psychophysics Toolbox Version 3 (Psychtoolbox-3).

#### Calibration of noise burst intensity

The intensity of the noise burst was calibrated using the method of constant stimuli. Specifically, ripples with noise burst intensities near and just above detection threshold (ranging from 0.0005 – 0.04 × intensity of the ripple in 11 steps) were generated and presented, along with ripple sounds without noise burst, in random order. Each of the 4 ripple sounds (2 center frequencies × 2 temporal modulation rates) was presented 12 times without a noise burst and repeated 4 times with each of the 11 burst sound intensities resulting in 56 trials per ripple. Participants were instructed to press a button if they heard a noise burst in the ripple sound. For each ripple center frequency, but combined across the two temporal modulation rates (112 trials in total), burst intensity was plotted against percentage hit (i.e., the percentage of trials on which the noise burst was correctly detected) and this relationship was fitted with a sigmoidal function. This allowed deriving the intensity at which the noise burst was detected with an accuracy of 60%, separately for 300 Hz and 4 kHz ripple sounds.

#### Behavioral data

Each participant completed a behavioral session that took place in a soundproof booth. In this session, we conducted the pure-tone audiogram and then calibrated the noise burst intensity. Next, each participant performed the noise burst detection task for the dual purpose of familiarizing the participants with the task before they entered the scanner and behaviorally validating the task. Specifically, participants were instructed to press a button whenever they heard a noise burst (whose intensity was set at 60% detection threshold) in the presented ripple sounds. Each of the four ripple sounds was presented 60 times per attentional condition, resulting in 240 trials per condition in total. These trials were divided into runs of 120 trials (i.e., two runs per condition). Inter-trial interval was equal to 1.7 s, resulting in a run duration of approximately 4 minutes (and approximately 16 minutes to complete this part of the behavioral data collection).

We analyzed the hit rate and d’ of all trials, as well as the reaction time of all trials that resulted in a hit (i.e., where the noise burst was correctly identified), with three separate two-way repeated measures ANOVAs (with factors “Sound Frequency” [300 Hz, 4 kHz] and “Condition” [Attend Low, Attend High]). Significant interactions were further explored through a paired t-test per level of the factor “Condition”.

#### MRI data

All measurements were performed on a 7 Tesla Siemens MAGNETOM scanner (Siemens Medical Solutions, Erlangen, Germany) using a single transmit 32 channel head coil (Nova Medical) at Scannexus (Maastricht, the Netherlands). Six of our eight participants took part in an earlier study (Sitek et al. 2019) consisting of three MRI measurement sessions (referred to below as MRI sessions 1-3). For these participants, only MRI sessions 4 and 5 were collected for this study and were added to the existing data. For the remaining two participants, data for all five MRI sessions was collected for this study.

In the first session, T_1_ weighted (T_1_w) and proton density weighted [PDw] data were collected at a voxel size of 0.7 mm isotropic. The T_1_w scan was acquired using a magnetization-prepared rapid gradient-echo (3-D MPRAGE) sequence (repetition time [TR] = 3100 ms; time to inversion [TI] = 1500 ms; time echo [TE] = 2.42 ms; flip angle = 5°; generalized autocalibrating partially parallel acquisition [GRAPPA] = 3; matrix size = 320 × 320; 256 slices). PDw images were acquired with the same 3-D MPRAGE as the T_1_w image, but without the inversion pulse (TR = 1380 ms; TE = 2.42 ms; flip angle = 5°; GRAPPA = 3; matrix size = 320 × 320; 256 slices; pixel bandwidth = 200 Hz/pixel). Dielectric pads were used to improve transmit efficiency in temporal areas when acquiring these anatomical images (Teeuwisse et al. 2012). Acquisition time for the T_1_w and PDw datasets were ~9.5 minutes each. In this first session, two additional anatomical datasets (a T_2_*-weighted and T_1_-weighted dataset with a short inversion time (Tourdias et al. 2014) and a diffusion-weighted MRI data were collected (see (Sitek et al. 2019) for acquisition parameters). These datasets were not used in this study.

In sessions 2-5, fMRI data were collected while participants listened to sounds. In sessions 2-3, 168 natural sounds (sound duration = 1 s) covering seven semantic categories (speech, voice, nature, tools, music, animals and monkey calls) were presented. Subjects performed a one-back task on the sounds, and pressed a button if consecutive sounds were repeated (trials with a repetition were excluded from the analysis). Following a rapid event-related design, sounds were presented in silent gaps in between fMRI acquisitions with an inter-trial interval of 2, 3, or 4 TRs. Session 2 and 3 were identical to each other. The 168 sounds were divided into four non-overlapping cross-validation sets of 42 sounds (each containing six samples per category). All sounds of a cross-validation set were presented once per run. As a session comprised 12 runs, each natural sound was repeated three times per session and six times throughout sessions 2-3 combined.

In sessions 4-5, participants listened to ripple sounds (sound duration = 1 s) with and without noise burst and performed the noise burst detection task. Noise burst intensity was set at 80%, as opposed to 60% for the behavioral data collected in the sound booth, as in pilot measurements it was observed that the task otherwise was too difficult to perform in the noisy scanner environment. The 80% noise burst detection threshold was determined by repeating a short version of the noise burst intensity calibration procedure inside the scanner. The procedure was kept the same as described above, except that each of the ripple sounds was presented 9 times (instead of 12) without a noise burst and repeated 3 times (instead of 4) with each of the 11 burst sound intensities resulting in 42 trials per ripple. The noise burst detection task was kept the same as in the behavioral part of the experiment as well. Participants had 1.7 s to respond to each sound. Upon button press, feedback was provided by a change in the color of the fixation cross (from black to green [correct] or red [incorrect]). Per run, each of the 4 ripples was presented 7 times (28 ripple trials/ run). Attentional condition was kept the same throughout a run. In “Attend Low” runs, 10 out of 14 ripples with a center frequency of 300 Hz contained a noise burst (71.4%), while the noise burst was present in only 4 out of 10 ripples with a center frequency of 4 kHz (28.6%). This was reversed in runs for the “Attend High” condition.

In addition to the ripple sounds, natural sounds (sound duration = 1 s, with 10 ms onset and offset linearly ramped) were presented interspersed with the ripple sounds for the purpose of estimating population receptive fields per attentional condition. Noise bursts did not occur in the natural sounds, and participants were informed accordingly. A total of 96 natural sounds was presented, comprising the following six sound categories: speech, voice, music, tools, animals, and nature sounds. The 96 natural sounds were divided into four non-overlapping cross-validation sets of 24 sounds (each containing 4 samples per category). Half of the sounds of a cross-validation set were presented once per run, and the other half was presented twice (resulting in 36 natural sound trials/ run). Across the two sessions, which comprised 16 runs (8 runs per session), each natural sound was repeated three times per attentional condition. Each run comprised 68 trials (28 ripples, 36 natural sounds, and 4 trials where no sound was presented). As all sounds were presented following the same rapid event related design as employed in sessions 2-3 (including the sound presentation during the silent gap between acquisitions), this resulted in a run duration just below 9 minutes. All sounds, both ripples and natural sounds, were sampled at 16 kHz. Sound energy was equalized using Matlab. Additionally, before onset of the scans but with earbuds in place, the loudness of the 300 Hz and 4 kHz ripple sounds was adjusted for each participant to match the loudness of the ripples to each other and to the loudness of the natural sounds. Sounds were presented to the participants in the MRI scanner using the MRI-compatible S14 model earbuds of the Sensimetrics Corporation (www.sens.com).

Functional MRI data throughout session 2-5 were acquired with a 2-D Multi-Band Echo Planar Imaging (2D-MB EPI) sequence (Moeller et al. 2010; Setsompop et al. 2012) (TR = 2600 ms; silent gap = 1400 ms; TE = 20 ms; flip angle = 80°; GRAPPA = 3; Multi-Band = 2; matrix size = 188 × 188; 46 slices; 1.1 mm isotropic voxels; phase encode direction inferior to superior). Acquisitions with reversed phase encode polarity were used for distortion correction. Slices were oriented coronally-oblique to cover the complete ascending auditory pathway (including the auditory brainstem structures, auditory thalamus, and auditory cortex).

#### MRI data analysis

The anatomical data analysis started by taking the ratio between the T_1_w and PDw images to minimize receive coil inhomogeneities in the T_1_w images (Van de Moortele et al. 2009). The resulting dataset was corrected for residual inhomogeneities, upsampled to 0.5 mm isotropic resolution, and brought to Talairach space. The white matter (WM) – gray matter (GM) boundary and the GM-cerebrospinal fluid (CSF) boundary were detected using the automatic tools of BrainVoyager QX, and then manually corrected. The WM-GM boundary was used for the cortical surface reconstruction of individual hemispheres, which were used for defining the following regions of interest based on macroanatomy (following the criteria outlined in Kim et al. (2000): core (Heschl’s gyrus), belt (planum temporale and planum polare), and parabelt (superior temporal gyrus). Furthermore, separate for the left and right hemisphere, we brought each hemisphere across participants to cortex-based aligned (CBA; Goebel et al., 2006) space for the purpose of group analysis.

We used BrainVoyager QX, FSL, and custom MATLAB code (The MATHWORKS Inc., Natick, MA, USA) to analyze the functional data. Preprocessing consisted of slice scan-time correction (with sinc interpolation), 3-dimensional motion correction, and temporal high pass filtering (6 sines/cosines). FSL-FLIRT was used to align the functional data of all sessions to those collected in the first run of session 2, while employing a mask that included the brainstem, thalamus and auditory cortex. The functional images were then distortion corrected using FSL-TOPUP based on the opposite phase encoding direction images collected in session 2. The functional data was the projected in Talairach space while re-sampling at a spatial resolution of 1 mm isotropic.

#### Responses to ripple sounds

We analyzed the functional data within a bilateral anatomical mask that covered the superior half of the temporal lobe, which includes the auditory cortex. A General Linear Model (GLM) analysis with a canonical hemodynamic response function (HRF) was used to estimate the overall BOLD response to the ripple and natural sounds in sessions 4-5, and thereby determine the auditory responsive regions in each participant. Next, we used a fixed-effects GLM analysis to evaluate the effect of selective attention on the BOLD responses to ripple sounds. In this second GLM analysis, we equalized the number of ripples with and without a noise burst across center frequencies within a run. That is, ripple sounds whose center frequencies matched the attended frequency were more often presented with noise burst (5 out of 7 ripples) than ripple sounds for which the center frequency did not match the attended frequency (2 out of 7 ripples). By excluding, per run, 3 ripples with(out) noise burst for ripples of the attended and non-attended frequency, respectively, we matched the number of presented noise bursts across ripple sounds. The effect of selective attention on the auditory cortical response to ripple sounds was first evaluated throughout the auditory cortex as a whole, by sampling the voxel timecourses to the reconstructed surfaces [spatial smoothing = 3 voxels], aligning them across participants in CBA space, and contrasting [attended ripple - non-attended ripple]. Second, we evaluated the effect of selective attention as a function of the difference between the attended frequency and the voxel’s best frequency [BF]. For this analysis, only voxels that showed a significant response to the sounds (FDR-corrected, *q* < 0.01) and a positive response (in percent signal change) to ripple sounds of both 300 Hz and 4 kHz in sessions 4-5 were included. We computed the change in response to each ripple sound when that sound was attended compared to non-attended as:

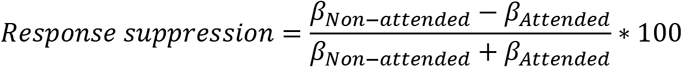

where β as the estimated response to the sound in percent signal change. Response suppression thus measures the effect of attention on the sound response, relative to the overall strength of the response. Response suppression was computed separately for voxels whose BF (averaged across attentional conditions) ranged from 0 to 1.5 octaves from the attended frequency (in 6 linearly-spaced bins). A linear regression analysis was used to test if response suppression (averaged across ripple sounds) varied with distance in octaves from the attended frequency (i.e., testing if the slope of the regression line was different from 0).

#### Population receptive field mapping

Responses to the natural sounds were used to estimate population receptive fields (pRFs) (Dumoulin and Wandell 2008; Thomas et al. 2015), separately for sessions 2-3 (“Baseline”) and the two attentional conditions of sessions 4-5 (“Attend Low” and “Attend High”). The response to the individual natural sounds was estimated following the procedure outlined in previous work (Moerel et al. 2012, 2015; Sitek et al. 2019). In short, separate for each cross-validation, GLM-denoise was used to denoise the data (http://kendrickkay.net/GLMdenoise/) (Kay et al. 2013). We then used a deconvolution GLM (nine stick predictors) to estimate the HRF separately for each voxel but common to all sounds (Kay et al. 2008). The resulting HRF and noise regressors, estimated on the training data, were used to compute a response estimate (beta weight) to the training and testing sounds in each voxel.

Following the methodology employed in pRF modeling of visual cortex (Dumoulin and Wandell 2008; Kay et al. 2015), we used a two step procedure to estimate the voxel-wise pRF based on its response estimate. First we estimated the best frequency and selectivity per voxel, and in a second step we estimated voxel gain. Through Fourier transform, we extracted the representation of each sound in the frequency space (logarithmically ranging from 180 to 8000 Hz in 2048 bins; Supplementary Figure S1A). The response to each sound was predicted by assuming a one dimensional Gaussian receptive field for each voxel, defined by the best frequency (mean μ of the Gaussian) and selectivity (size σ of the Gaussian). The possible parameter ranges of the one dimensional Gaussian were defined by creating a regular grid of 205 seeds in the frequency space (logarithmically ranging from 180 to 8000 Hz) and 10 seeds in the selectivity space (ranging from 0.27 to 2.67 octaves; Supplementary Figure S1). For each point in this grid, a predicted sound response (in beta weights) was generated by multiplying the sound representation in frequency space with the frequency response curve defined by that grid point. For each voxel and gridpoint, we then computed a correlation coefficient between the observed and predicted beta responses.

As we have shown before (Lage-Castellanos et al. 2020), the distribution of the resulting cost function under the null hypothesis varies with the pRF selectivity. As a result, the gridpoint generating the most correlated prediction with the observed data is not necessarily the least probable one to occur by chance. We therefore followed the Permutation Based Model Grid-Search for Separable Betas Design (PermGS) procedure, described in detail in (Lage-Castellanos et al. 2020). This is a refinement of the grid search pRF-estimation procedure where the selection criterion varies from the traditional cost function (correlation coefficient) to a criterion based on the probability of selecting a particular seed under the null (−log[p value]). While similar to the standard grid search algorithm, this variant prevents the bias towards high selectivity pRFs (small pRF size) in voxels with low SNR. A total of 1000 permutations were implemented, where in each permutation the sound order (i.e., the vector of beta responses) was randomized. The distribution of the correlation coefficient between observed and predicted beta response was computed for each seed at each randomization. The best seed for the observed data was selected as the one with the lowest probability of occurrence under the null hypothesis across all the null distributions for each seed in the grid. As shown in (Lage-Castellanos et al. 2020), this change in the selection criterion does not modify the pRF parameters in voxels with a good SNR, but increases the reliability of the estimated pRFs in voxels with a low SNR.

The pRF fitting procedure was performed per dataset (separate for Baseline, Attend Low, and Attend High). Each dataset was designed to contain four non-overlapping sets of sounds presented in separate runs. Fitting was performed on training data in 4-fold cross validation, where each training dataset consisted of 3 out of 4 sound sets. Prediction accuracy was assessed by correlating the predicted and observed voxels’ responses in independent testing data (i.e., the left out sound set). The final prediction accuracy was computed as the average accuracy on the testing data across the four folds.

After selecting the best frequency μ and selectivity σ per voxel, we estimated the voxel’s gain (in percent signal change [PSC]) per attentional condition as the beta weight resulting from the linear regression between the observed and predicted voxel’s time series. In this regression analysis, the predicted voxel’s time series was based on the frequency content of the presented sound, the attentional condition, and the condition- and voxel-specific pRF estimate. Noise covariates (as estimated through GLM-denoise) were included as predictors in the regression analysis. Gain was computed in 4-fold cross validation.

The cortical maps of frequency preference and selectivity were created by color-coding each voxel according to the mean (best frequency [BF]) and size (tuning width [TW]) of the best fitting Gaussian, averaged over cross-validations. A red-yellow-green-blue color scale was used to create BF maps, where preference for low and high frequencies was assigned to red and blue colors, respectively. A yellow-green-blue-purple color scale was used for the TW maps, where broad and narrow TW were assigned with yellow and purple colors, respectively. Maps were restricted to those voxels that showed a significant response to the sounds (FDR-corrected, *q* < 0.01) in sessions 4-5. Group maps were created by sampling the individual maps to CBA space, smoothing each map at individual participant level (FWHM = 2.4 mm), and averaging the resulting maps at all locations for which data of at least 5 out of 8 participants was available. To evaluate the existence of a relation between maps of BF and TW, we computed the Pearson’s correlation between these maps in each individual participant and per attentional condition. Per attentional condition, a one sample t-test was used to test the statistical significance of the observed Pearson’s correlation between BF and TW maps across the 8 participants.

#### Evaluation of attention-induced pRF changes

In addition to limiting the analysis to those voxels with a significant response to the sounds, for all following analyses we furthermore limited the voxels to those that in the Baseline dataset (scanning sessions 2-3; averaged across the four cross-validations) showed a Pearson correlation between predicted and observed responses to testing sounds > 0.18 (which corresponds to *p* < 0.01, determined through permutation testing). Importantly, the Attend Low and Attend High datasets (scanning sessions 4-5) were not used for voxel selection. We reasoned that if pRFs changed across attentional conditions, the prediction of responses to test sounds should be more accurate if based on data of the same condition (within-condition prediction accuracy) than on data of a different condition (across-condition prediction accuracy). To test for the presence of pRF changes with attention, we therefore computed four maps of prediction accuracy per participant, where pRFs were computed on either training dataset (“Attend Low” or “Attend High”) and evaluated on either testing dataset (“Attend Low” or “Attend High”). We then subtracted maps of within-condition prediction accuracy from across-condition prediction accuracy (i.e., “Low-to-Low” – “High-to-Low”, and “High-to-High” – “Low-to-High”), projected these individual maps to the surface and brought them to CBA space, and averaged the maps across conditions and participants. Cluster-size permutation thresholding, performed separately for the left and right hemisphere, was used to determine the statistical significance of the activation clusters observed in the resulting group maps. Specifically, by performing all possible sign inversions of the 8 participant maps (2^8^ = 256 permutations), thresholding the maps (at 7 different thresholds that ranged from 0.03 – 0.09 in linearly-spaced bins) and recording the maximum cluster sizes occurring for each permuted group map (using the SurfStat toolbox; https://www.math.mcgill.ca/keith/surfstat/), we constructed the null distribution of the maximum cluster size. The 95^th^ percentile of this distribution was used as the cluster size threshold for determining significance of clusters in the observed data. This imposes a family wise error rate (FWER) control at the level of 0.05.

We visualized the distribution of BF, TW, and gain, separately for the core (Heschl’s gyrus), belt (planum temporale and planum polare), and parabelt (superior temporal gyrus), through histograms (10 linearly-spaced bins). These histograms were computed per participant, and normalized for the total number of included voxels at individual subject level. Per region of interest, paired t-tests were used to test for differences in average BF, TW, and gain across attentional conditions.

We also evaluated the effect of selective attention on TW and gain as a function of the difference between the attended frequency and the voxel’s best frequency [BF]. We computed the average TW and gain in bins relative to the attended frequency (ranging from 0 – 1.5 octave in 6 linearly-spaced bins), separately for when that frequency was attended and was not attended. Separately for TW and gain, a two-way repeated measures ANOVA per region of interest with factors ‘Attention’ (2 levels: attended vs. non-attended) and ‘Distance to target frequency’ (6 levels: 0 - 0.25 octaves, 0.25 - 0.5 octaves, 0.5 - 0.75 octaves, 0.75 - 1 octaves, 1 – 1.25 octaves, and 1.25 – 1.5 octaves) was used to test for differences in with selective attention. Significant interactions were further explored through a paired t-test per level of the factor ‘Distance to target frequency’.

## Results

Selective attention sped up the reaction time to noise bursts in attended compared to unattended ripples (Figure 1A-B). This effect was significant when probed outside the scanner (Fig. 1A; significant interaction ‘Sound Frequency’ × ‘Condition’; F(1,6) = 11.73; *p* = 0.011). Follow up tests per level of ‘Sound Frequency’ showed a significantly faster response to noise bursts in 300 Hz ripples when attended (Bonferroni corrected *p* = 0.002). The reaction time to noise bursts in 4 kHz ripple sounds did not significantly differ between attentional conditions. While inside the scanner (i.e., during MRI data collection) reaction times qualitatively followed the same pattern as in the soundproof booth (Fig. 1B), we did not observe a significant interaction between ‘Sound Frequency’ and ‘Condition’ nor any significant main effects.

**Figure 1.**
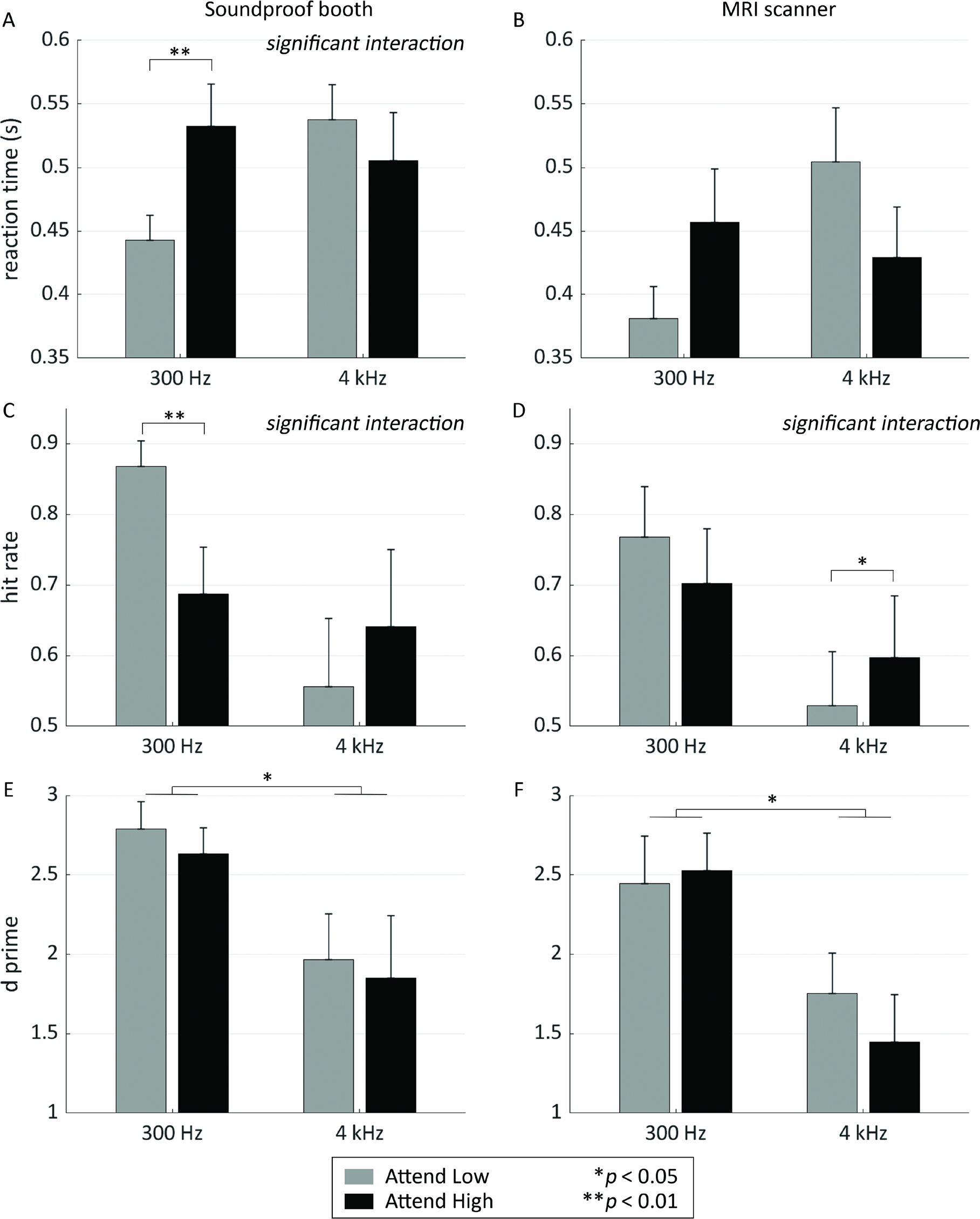
Effect of attention on behavioral responses. Selective attention sped up the reaction time to noise bursts that occurred in ripples of the attended frequency. This effect was present when probed outside and inside the scanner (A-B, respectively), but only significant when tested outside the scanner. C-D) Hit rate increased with attention, both outside and inside the scanner (in the left and right panel, respectively). E-F) Attention did not affect detectability index d’, but we did observe an overall higher detectability for low compared to high frequency ripples. This pattern was present both outside and inside the scanner (in the left and right panel, respectively). Throughout figure panels, error bars show the standard error across participants. Single and double asterisks indicate statistically significant differences between conditions at *p* < 0.05 and *p* < 0.01, respectively.

We also observed a facilitatory effect of selective attention on the hit rate (sound booth: significant interaction, F(1,6) = 32.78; *p* = 7.2 × 10^−4^; significant difference between levels of ‘Condition’ for 300 Hz sounds, Bonferroni corrected *p* = 0.0063; inside the scanner: significant interaction, F(1,6) = 8.53; *p* = 0.022; significant difference between levels of ‘Condition’ for 4 kHz sounds, Bonferroni corrected *p* = 0.040; Fig. 1C-D). However, this was accompanied by an increase in the number of false alarms for attended ripples and therefore likely due to a response bias. Indeed, we did not observe an effect of selective attention on d’ (Fig. 1E-F). There was, however, a main effect of ‘Sound Frequency’ on d’ (sound booth: F(1,6) = 7.51; *p* = 0.029; inside the scanner: F(1,6) = 6.01; *p* = 0.044), indicating a higher detectability of noise bursts in ripples of 300 Hz compared to 4 kHz in spite of the noise burst intensity calibration.

We observed significant responses (FDR-corrected, *q* < 0.05) to the sounds throughout the auditory cortex. The responses to ripple sounds were weaker when these sounds were attended compared to unattended. This effect was observed throughout the auditory cortex (Figure 2A), especially in the belt and parabelt regions of the right hemisphere. In order to further examine the weaker response to attended sounds, we assessed the voxels’ frequency preference (best frequency; BF), frequency selectivity (tuning width; TW), and gain through pRF mapping based on responses to the natural sounds. We estimated separate pRF’s for the three conditions (“Baseline”, “Attend Low” condition, and “Attend High”). PRF’s could be reliably estimated throughout individual participants and conditions (see Supplementary Figure S2 for pRF fits in example voxels), showing a prediction accuracy in testing that was consistently above zero (see Figure 3 for within-condition prediction accuracy, and Supplementary Figure S3 for across-condition prediction accuracy). When analyzing attention-induced suppression of responses to ripple sounds as a function of distance between the voxel’s BF and the attended sound frequency, we observed a stronger suppression in those voxels whose BF most closely matched the attended sound frequency. Response suppression for both ripple frequencies decreased with increasing distance between the voxels’ BF and the attended sound frequency (Figure 2B). Statistical testing on the average response suppression across ripple frequencies showed a significant linear trend (*p* = 0.011).

**Figure 2.**
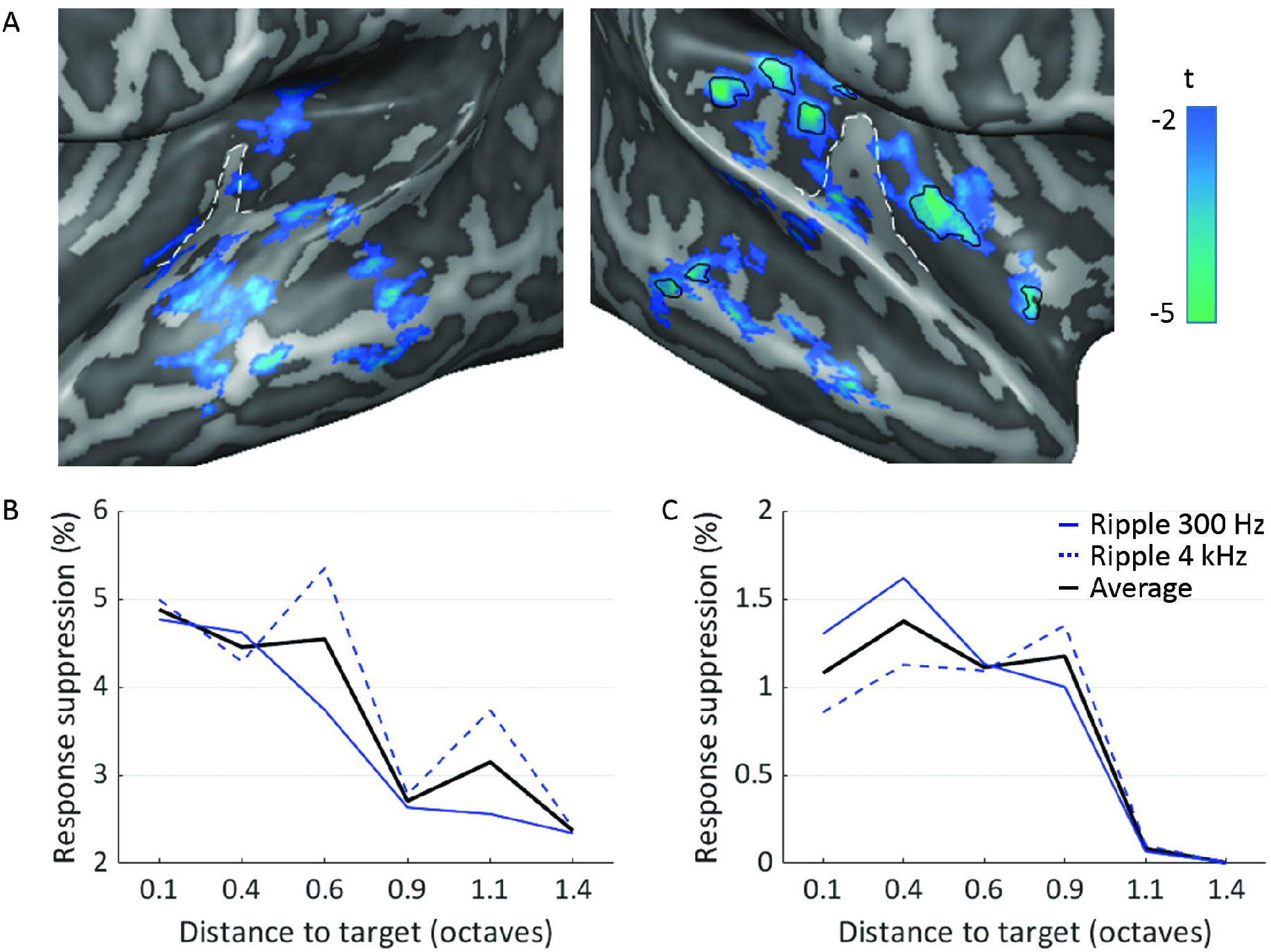
Effect of selective attention on response to ripples. A) At group level, the response to attended ripple sounds was lower than when the same sounds were non-attended ([response to attended – response to non-attended sounds] shown in blue-green). This effect could be observed throughout the auditory cortex (*p* < 0.05 uncorrected), but was strongest in belt and parabelt regions of the right hemisphere (FDR-corrected, *q* < 0.05 shown in black outlines). The white dashed line outlines Heschl’s gyrus (HG). B) Response suppression (in %) to ripple sounds when attended compared to non-attended as a function of the distance (in octaves) between the voxel’s BF and the attended (i.e., target) frequency. Suppression of responses to ripple sounds at 300 Hz (solid blue line), 4 kHz (dashed blue line), and their average is shown. Responses to attended ripples were suppressed compared to when the same ripple sounds were not attended, and this effect was stronger in voxels with a BF that closely matched the attended frequency. C) Modeled response suppression (%) to attended compared to non-attended ripple sounds as a function of the distance (in octaves) between the voxels BF and the attended (i.e., target) frequency. Modeled response suppression to ripple sounds at 300 Hz (solid blue line), 4 kHz (dashed blue line), and their average is shown. Model responses to attended ripples in voxels with a BF that closely matched the attended frequency were weaker than when the same ripple sounds were not attended.

**Figure 3.**
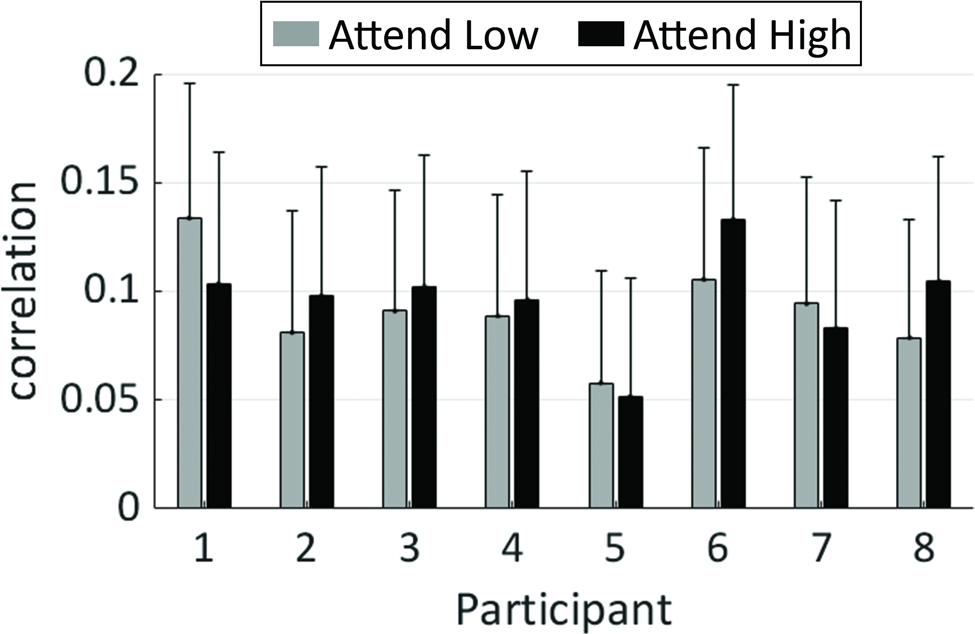
Prediction accuracy. Prediction accuracy as the Pearson correlation coefficient between predicted and observed responses to test sounds, averaged across voxels and cross validations, in each participant. All voxels with significant response to sounds and a prediction accuracy > 0.18 (corresponding to *p* < 0.01) in the “Baseline” dataset were included. Gray and black bars show results from the “Attend Low” and “Attend High” condition, respectively. Error bars represent the standard deviation across cross validations.

To examine the processing changes that may have caused this weaker response to the attended ripple sounds, we next explored the voxels frequency preference, frequency selectivity and gain, as well as changes in voxel tuning with attention. The BF maps (i.e., tonotopic maps) were in accordance with previous reports (Humphries et al. 2010; Da Costa et al. 2011; Striem-Amit et al. 2011; Langers and van Dijk 2012; Moerel et al. 2012). Prediction accuracy varied with BF, such that voxels with a BF around 2 kHz had the lowest prediction accuracy (Supplementary Figure S3). The lower prediction accuracy in the middle frequency range likely resulted from its coincidence with the spectrum of the scanner noise. Maps of tuning width (TW) followed previous reports as well (Kajikawa 2004; Rauschecker and Tian 2004; Kusmierek and Rauschecker 2009; Moerel et al. 2012) and showed high correspondence across conditions (see Figure 4 for maps of individual participant S01 and for the group maps). In accordance with previous findings (Cheung et al. 2001; Imaizumi et al. 2004; Moerel et al. 2012), BF (in Hz) was negatively correlated with tuning width (TW, in octaves), such that a higher BF corresponded to more narrow tuning in the “Baseline” maps (mean [SE] correlation across subjects = −0.07 [0.02]; one-sample t-test, t(7) = −3.16, *p* = 0.016). For maps in conditions “Attend Low” and “Attend High” we did not observe a significant correlation between BF and TW (mean [SE] correlation across subjects = −0.03 [0.02]; one-sample t-test, t(7) = −1.40, *p* = 0.203 for “Attend Low”, and mean [SE] correlation across subjects = −0.04 [0.02]; one-sample t-test, t(7) = −2.24, *p* = 0.060 for “Attend High”).

**Figure 4.**
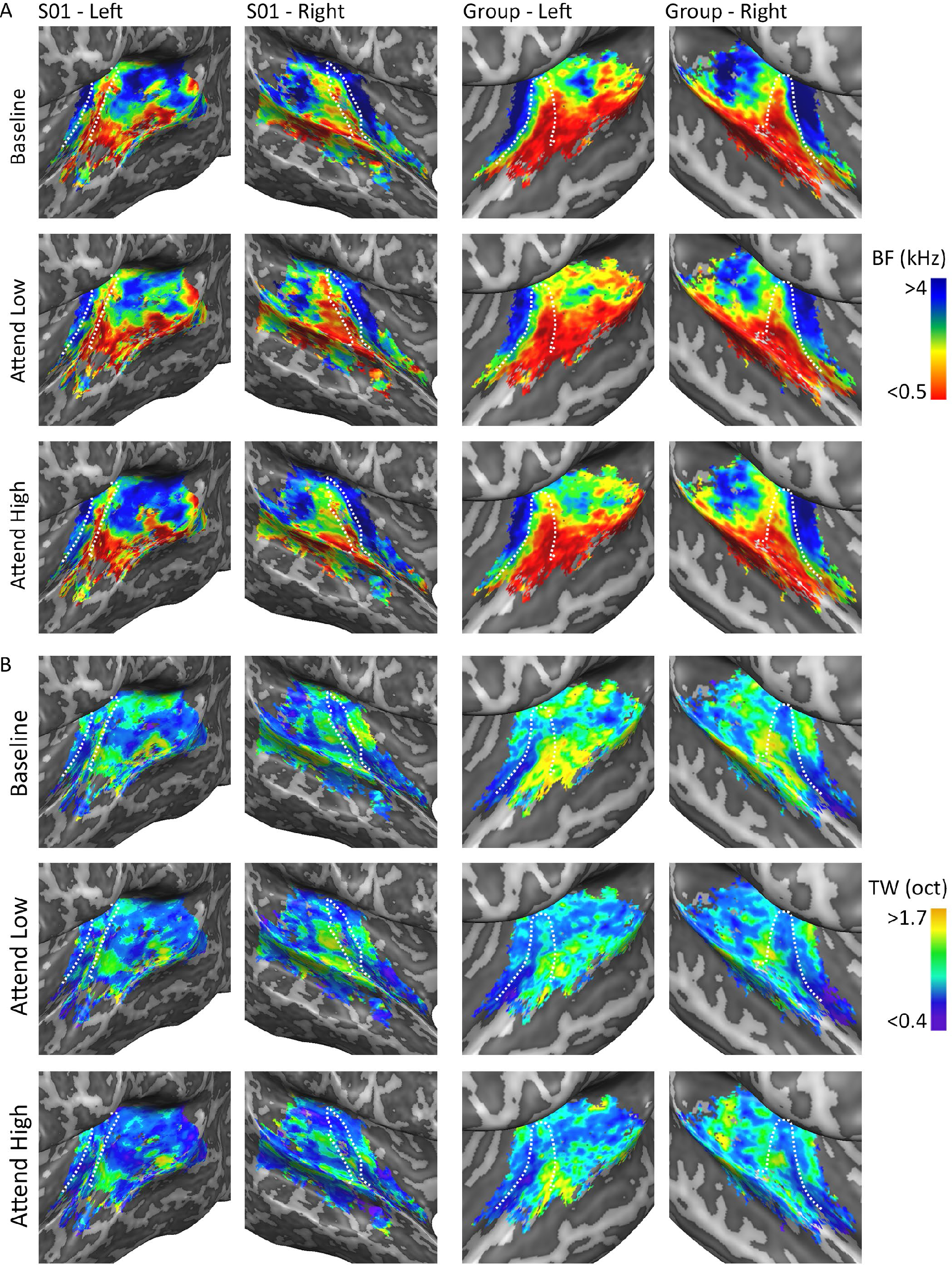
Maps of frequency preference and selectivity. Tonotopic maps (i.e., best frequency [BF]) and maps of tuning width (TW) per hemisphere, in a single participant (S01; left) and at group level (right). Maps showed high correspondence across conditions. The white dashed line outlines HG.

As a next step, we examined if pRFs changed across attentional conditions. We reasoned that if pRFs changed, the prediction accuracy when training and testing on natural sound responses originating from within the same attentional condition should be higher than if training and testing sounds originated from different attentional conditions. Indeed, group maps of cross modal difference (computed as [within-condition accuracy – across-condition accuracy]) were overall positive, indicating higher within-condition prediction accuracy than across-condition prediction accuracy (Figure 5A). Cluster-size permutation thresholding showed that the size of the positive clusters in the map (i.e., where within-condition accuracy > across-condition accuracy) across a range of map thresholds were greater than expected by chance (Figure 5B). This was not the case for the negative clusters in the map (i.e., where within-condition accuracy < across-condition accuracy; Figure 5B). The same results were observed when analyzing the cross modal differences separately per condition (i.e., when model training was either performed on data of the Attend Low condition or on data of the Attend High condition; Supplementary Figure S4). These results confirm that in parts of the auditory cortex the within-condition prediction accuracy was indeed greater than the across-condition accuracy, indicating the presence of pRF changes across attentional conditions.

**Figure 5.**
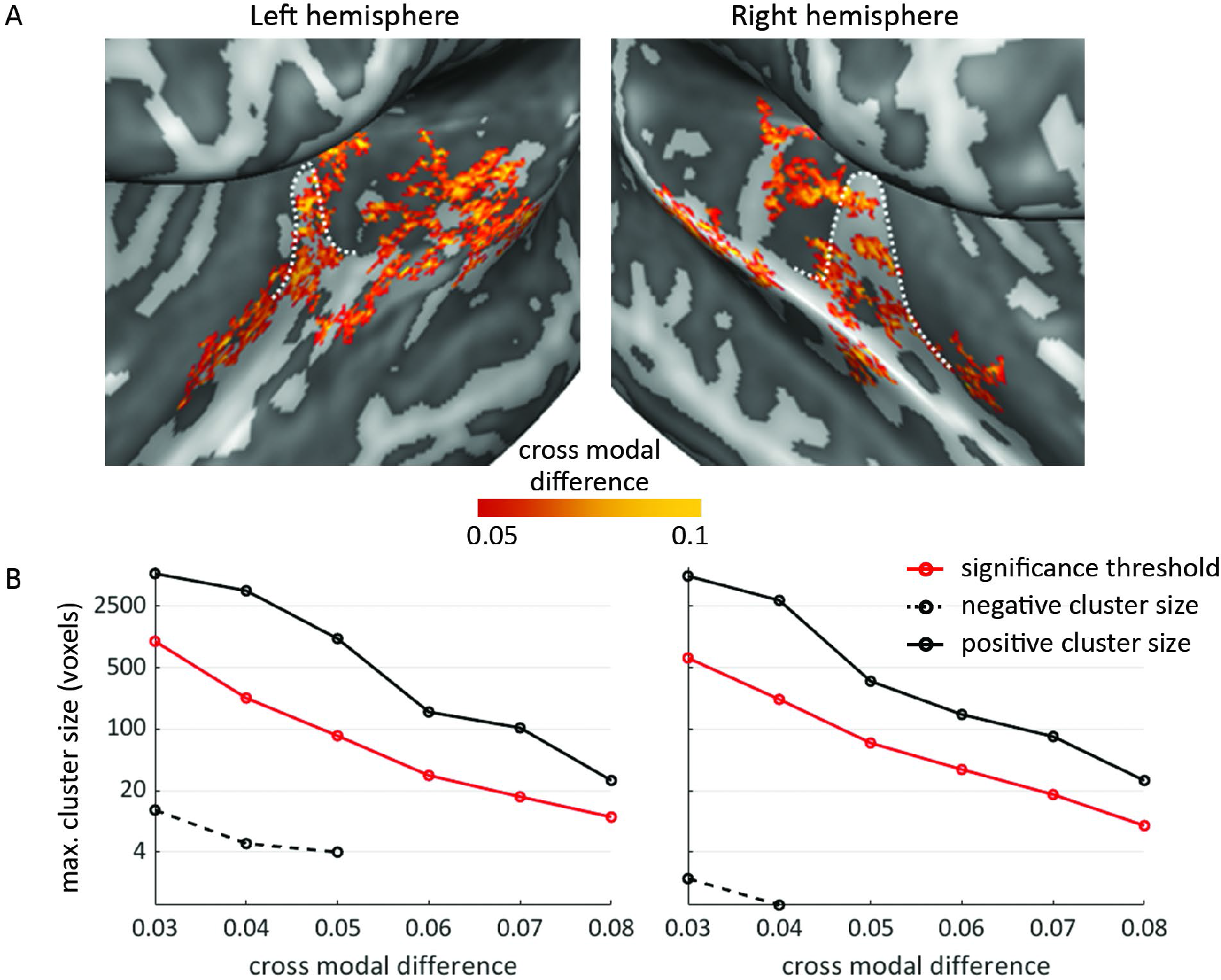
Within- compared to across-condition prediction accuracy. A) The cross modal difference (computed as [within-condition – across-condition prediction accuracy]) maps show positive values, indicating that prediction accuracy was higher when probed within- compared to across-attentional conditions. The white dashed line outlines HG. Maps are cluster size corrected at a FWER = 0.05. B) Across hemispheres, the largest positive cluster size observed (solid black line) was greater than expected by chance (red line). The largest negative cluster size observed (dashed black line) did not exceed the cluster size expected by chance (red line). This was true independent of the “cross modal difference” threshold. Note that for higher “cross modal difference” thresholds, no negative clusters were present in the maps and correspondingly no dashed black line is shown at these thresholds.

To examine pRF changes with attention, we created BF, TW, and gain histograms across conditions, separately for the core, belt, and parabelt (see Supplementary Figure S5 for regions of interest) for all pRFs in these regions. No major differences between conditions in BF, TW, or gain were apparent (Figure 6A-C). Indeed, when comparing the pRF parameters across the two attentional conditions, we did not observe a significant difference in any of the regions of interest (core: two-sided paired t-test on BF: t(7) = −0.134, *p* = 0.897; two-sided paired t-test on TW: t(7) = 1.046, *p* = 0.331; two-sided paired t-test on gain: t(7) = −0.260, *p* = 0.802; belt: two-sided paired t-test on BF: t(7) = −0.795, *p* = 0.453; two-sided paired t-test on TW: t(7) = −1.128, *p* = 0.297; two-sided paired t-test on gain: t(7) = −0.163, *p* = 0.875; parabelt: two-sided paired t-test on BF: t(7) = 0.747, *p* = 0.479; two-sided paired t-test on TW: t(7) = −0.584, *p* = 0.578; two-sided paired t-test on gain: t(7) = −0.315, *p* = 0.762.

**Figure 6.**
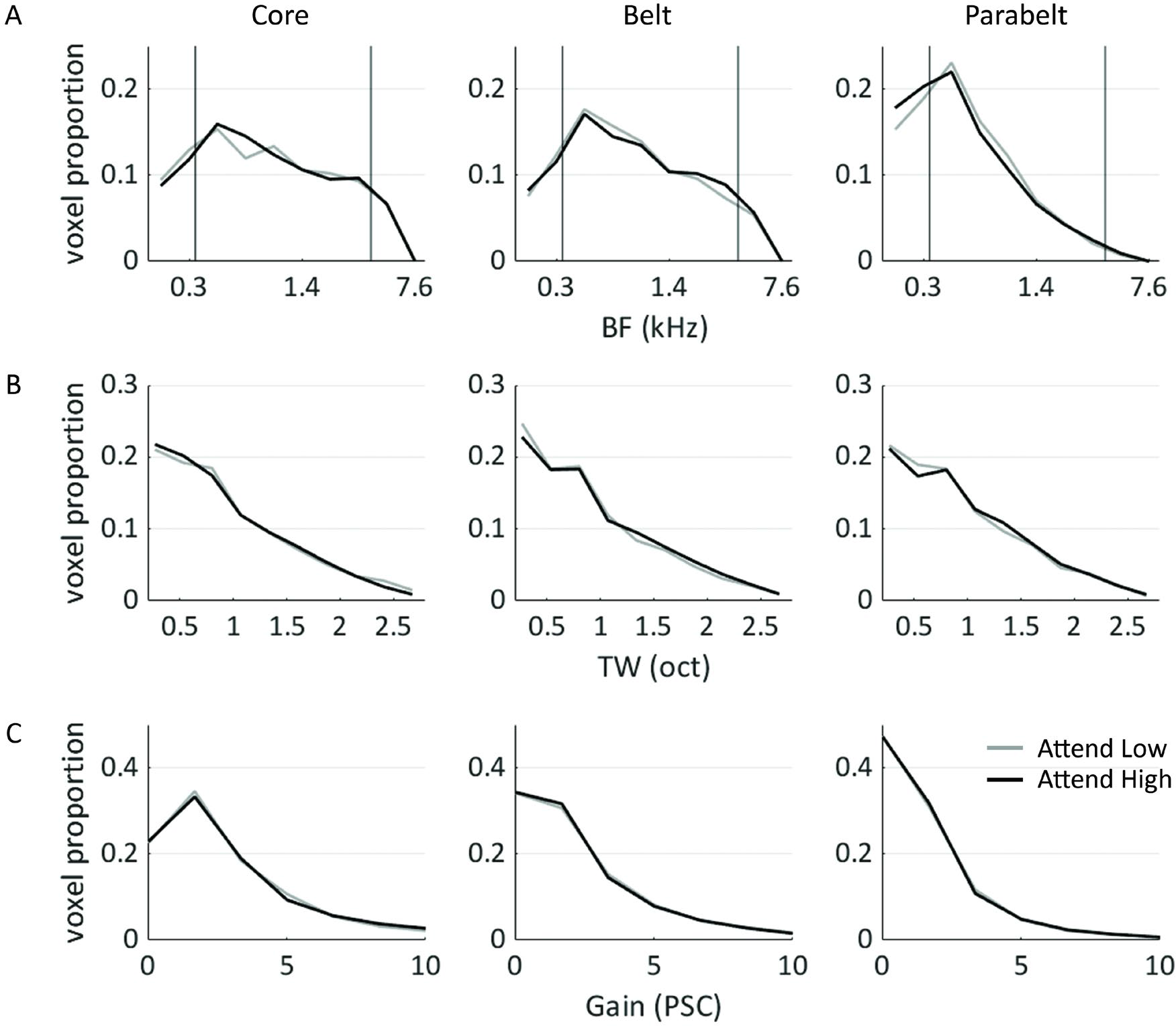
Distribution of frequency preference, frequency selectivity, and gain. Distribution of frequency preference (BF, in kHz), frequency selectivity (TW, in octaves), and gain (in percent signal change [PSC]) are shown from top to bottom, across core, belt, and parabelt shown from left to right. In A, the attended frequencies are indicated by black vertical lines. No differences between “Attend Low” (gray line) and “Attend High” (black line) were apparent.

Following our observation that the largest changes in ripple responses occurred in voxels whose BF was close to the attended frequency, we then explored TW and gain changes as a function of voxel BF. This analysis showed a TW change with attention in the auditory parabelt. Specifically, while no significant attention-induced TW differences were observed in the core (two-way repeated measures ANOVA with factors ‘Attention’ and ‘Distance to target frequency’; no significant interaction or main effect of ‘Attention’; significant main effect of ‘Distance to target frequency’, F(1,5) = 35.426; *p* = 9.57 × 10^−13^) or belt (two-way repeated measures ANOVA with factors ‘Attention’ and ‘Distance to target frequency’; no significant interaction or main effect of ‘Attention’; significant main effect of ‘Distance to target frequency’, F(1,5) = 41.591; *p* = 9.23 × 10^−14^), TW narrowed in parabelt voxels with attention (Figure 7A). This effect was not driven by a sampling bias, as TW narrowing in the auditory parabelt was observed both when the target frequency was 300 Hz and 4 kHz (Supplementary Figure S6). The effect of attention on TW was stronger in voxels with a BF close to the attended frequency (significant interaction between factors ‘Attention’ and ‘Distance to target frequency’ (F(1,5) = 3.522; *p* = 0.011), specifically in voxels with a BF up to 0.5 octaves distance from the attended frequency (follow-up paired t-tests per level of factor ‘Distance to target frequency’; significant difference in bin 0-0.25 octaves before but not after multiple comparison correction [t = −2.606; *p_uncorr_* = 0.018; *p_corr_* = 0.105; significant difference in bin 0.25 – 0.5 octaves [t = −3.669; *p_uncorr_* = 0.004; *p_corr_* = 0.024; no trends towards significant differences in the other bins). No significant attention-induced gain changes were observed in the core (two-way repeated measures ANOVA with factors ‘Attention’ and ‘Distance to target frequency’; no significant interaction or main effect of ‘Attention’; significant main effect of ‘Distance to target frequency’, F(1,5) = 3.414; *p* = 0.013), belt (two-way repeated measures ANOVA with factors ‘Attention’ and ‘Distance to target frequency’; no significant interaction or main effect of ‘Attention’; significant main effect of ‘Distance to target frequency’, F(1,5) = 13.192; *p* = 2.99 × 10^−7^), or parabelt (two-way repeated measures ANOVA with factors ‘Attention’ and ‘Distance to target frequency’; no significant interaction, main effect of ‘Attention’ or ‘Distance to target frequency’; Figure 7B).

**Figure 7.**
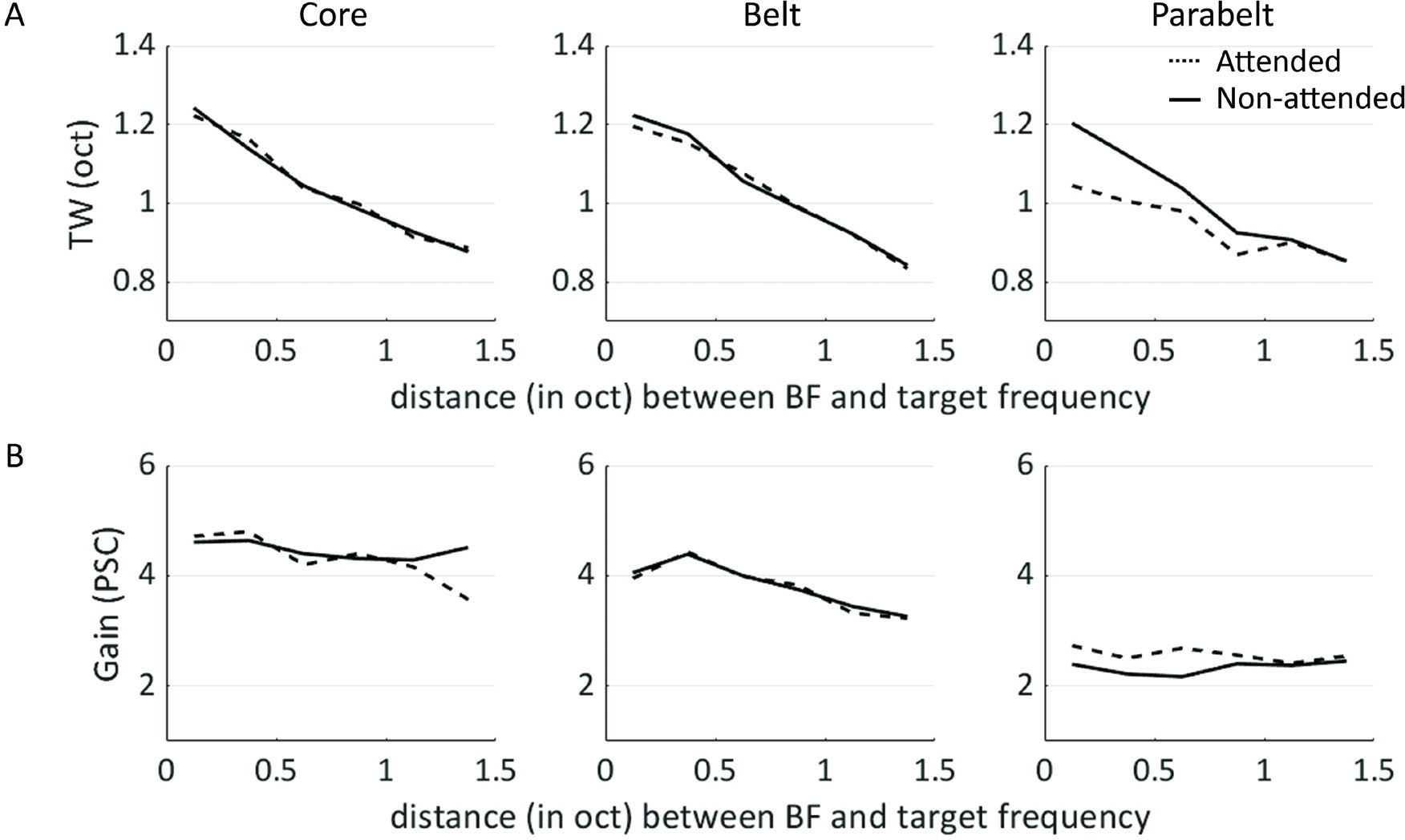
Effect of attention on frequency selectivity and gain. A) Tuning width (TW) as a function of distance (in octaves) between BF and the attended target frequency. In the parabelt, TW was more narrow when the BF was attended (dashed black line) compared to when it was not attended (solid black line). This effect was strongest in those voxels whose BF was closest to the attended sound frequency. B) Gain as a function of distance (in octaves) between BF and the attended target frequency. No attention-dependent differences in gain were observed.

As a final step, we used computational modeling to explore how the pRF changes observed in our data related to the observed fMRI responses to ripple sounds. To this end, we modeled the auditory cortex as a set of voxels whose response was fully characterized by their frequency preference and selectivity. BF was set to the frequency preference (averaged over attentional conditions) observed in the left hemisphere of a representative participant (S01 reported in Fig. 4A). While BF was the same across attentional conditions, TW was modelled to sharpen with attention in voxels with a BF close to the attended one. TW parameters across attentional conditions were set to match the results reported in Fig. 7A. Gain was kept the same across voxels, as no attention-dependent effect on gain was observed. Responses to ripple sounds per attentional condition were modeled as the multiplication of their sound spectrum with the condition-specific pRF of each voxel and normalized between 1 and 2 percent signal change to match the response strength observed in our dataset. We then explored TW changes as a function of the difference between the attended frequency and the voxel’s best frequency [BF] following the same procedure as used on our data. That is, we computed response suppression with attention in voxel bins defined relative to the target frequency (ranging from 0 – 1.5 octave in 6 linearly-spaced bins). In accordance with our data (reported in Fig. 2B), modeled responses to attended ripples in voxels with a BF that closely matched the attended frequency were weaker than when the same ripple sounds were not attended (Fig. 2C). While model output suggested that tuning width narrowing could account for approximately 1.5% suppression in responses to ripple sounds, our data showed up to 5% response suppression (compare panels B-C in Figure 2). This indicates that the observed weaker response to attended ripple sounds in our dataset could have been at least in part driven by pRF narrowing.

## Discussion

Our selective attention task resulted in a faster response to targets in attended compared to non-attended sounds. Task performance was accompanied by a lower response in auditory cortex to attended sounds, especially in voxels whose frequency preference closely matched the attended sound frequency. This observation was surprising, as previous fMRI studies of human auditory cortex reported increased BOLD responses, interpreted as increased gain, with attention (Paltoglou et al. 2009; Da Costa et al. 2013a; Dick et al. 2017; Riecke et al. 2017, 2018). Our results refute increased gain at voxel level as a model of selective attention in our task.

A response reduction with attention is in line with results of previous animal electrophysiology studies (David et al. 2012). As these studies also reported receptive field changes with attention (Fritz et al. 2003, 2007; Lee and Middlebrooks 2011), we next examined if we could shed light on the observed attention-induced response reduction through population receptive field mapping. With this approach, we specifically explored changes in frequency preference, frequency selectivity, and gain with attention. Frequency preference and gain were not modulated by attention. Instead, frequency selectivity (tuning width) differed across attentional conditions. That is, we observed narrower frequency tuning with attention in auditory cortical parabelt locations with a preferred frequency close to the attended one. Based on responses of a simple computational model of auditory cortex, we suggest that the weaker response to attended ripple sounds in our dataset was at least in part driven by pRF narrowing.

Model output suggested that tuning width narrowing could account for approximately 1.5% suppression in responses to ripple sounds. Instead, our data showed up to 5% response suppression. The simplicity of and assumptions behind our modeling approach could be responsible for this discrepancy. Estimation of feature tuning based on fMRI data can be performed in multiple ways. We characterized the voxel’s population receptive field by a one dimensional Gaussian function. This represents a simplification of voxel tuning, as a one dimensional Gaussian cannot characterize complex, multi-peaked frequency preferences that are known to be present in primary and especially higher order auditory regions (Kadia 2002; Sadagopan and Wang 2009; Moerel et al. 2013; Kikuchi et al. 2014). It is possible that we failed to pick up attention-induced changes beyond the main frequency preference, and that these changes account for the part of the observed responses reduction that currently remains unexplained. The employment of fMRI encoding techniques (Moerel et al. 2013; Santoro et al. 2014) as opposed to the population receptive field mapping would have allowed assessment of complex, multi-peaked voxel tuning. However, compared to fMRI encoding, our current approach is preferential for reliably assessing attention-induced changes in the voxel’s population receptive field. While estimation of frequency preference (BF) and selectivity (TW) based on encoding is possible, estimation of these parameters has been shown to be more reliable through pRF mapping (Lage-Castellanos et al. 2020). Moreover, in presence of regularization (which is the standard approach in encoding) (Moerel et al. 2013; Santoro et al. 2014) the comparison of resulting parameters (i.e., the pRF) across conditions is not straightforward due to possible differences in regularization strength across conditions. Indeed, population receptive field mapping as employed here was previously shown to be reliable for characterizing auditory tuning (Thomas et al. 2015) as well as for assessing the effect of spatial attention along the ventral temporal cortex in the visual domain (Kay et al. 2015).

While previous fMRI studies reported increased gain with attention, we instead report a change in voxel tuning. What caused the striking difference between our results compared to previous fMRI studies? We would like to propose two tentative hypotheses to serve as the basis for follow up research. First, the difference in results may have originated from the difference in experimental design. Previous fMRI studies simultaneously presented the target (attended) and reference (non-attended) sounds in a scene, while our stimulus presentation was sequential (i.e., consecutive presentation of target and reference sounds). While simultaneous presentation requires enhancing attended sounds compared to background noise, such a strategy may not be needed when only one sound stream is present. In sequential presentation, inhibiting responses to irrelevant stimuli (as operationalized through increased frequency selectivity) may be more beneficial than increasing responses to relevant ones. In line with findings from electrophysiological studies in animal models highlighting the diversity in the manner by which attention influences auditory cortical processing (Atiani et al. 2009; David et al. 2012), our results suggest that fundamentally different mechanisms may underlie auditory selective attention when sounds are presented in a scene compared to when sounds are presented consecutively. A second, alternative hypothesis for explaining the difference between our results and those of previous fMRI studies is that various attentional mechanisms may differentially dominate the hierarchical processing stages. In early auditory cortex, where neuronal (and voxel) tuning is rather simple, a gain mechanism may be sufficient to enhance responses to attended acoustic features and reduce responses to non-attended features. Instead, such a mechanism may be insufficient for higher order auditory cortex, as both attended and non-attended acoustic features may fall within a neuron’s broad and complex tuning profile. Here, simplification of neuronal tuning – for example through a narrowing of tuning width – could be a computationally superior attentional mechanism. Indeed, previous invasive animal electrophysiology studies reported an increase in attention-induced retuning at higher levels of the auditory processing hierarchy (Atiani et al. 2014; Elgueda et al. 2019). This hypothesis is also in line with results from a multimodal M/EEG and fMRI study showing attention-driven retuning of neural responses, but not gain, in human non-primary auditory cortex (Ahveninen et al. 2011).

In addition to the attention-induced narrowing of frequency tuning, we observed an overall (i.e., attention-independent) sharpening of tuning width with increasing distance between voxel’s preferred frequency and the attended frequency. This could reflect a true neurobiological finding, i.e., a condition-independent narrowing of tuning width for the middle frequency range. Alternatively, the sharpening of tuning width could be an artifact of the pRF estimation. The accuracy of pRF estimation was lowest for voxels in the middle frequency range (~2 kHz; Supplementary Figure S3B). The lower prediction accuracy in the 2 kHz range is in accordance with our previous observations (Santoro et al. 2017) and likely results from the fMRI noise. That is, the BOLD response may be partially saturated in those neuronal population whose frequency preference matches the spectrum of the scanner noise. PRF estimation is biased towards selecting more narrowly tuned Gaussian profiles in cases of low SNR (Lage-Castellanos et al. 2020). While the pRF estimation approach was optimized to address this bias, we cannot exclude that it was still partially present for voxels with the lowest SNR. As the affected frequency region is far away from the attended regions, we do not expect this to affect our main conclusions.

In conclusion, our results argue for the existence of diverse attention-induced effects in auditory cortex that may depend on task, setting, and hierarchical auditory processing stage. These observations urge for future work that directly compares tasks with regard to their influence on sensory representation throughout auditory cortex. It would furthermore be of interest to explore the influence of the stimulus characteristics embedding the attended feature, and the effect of switching features (e.g., attending specific spectral or temporal modulations instead of a specific frequency). Furthermore, studying the relationship between which feature is attended and how stably this feature is encoding throughout cortical depth (O’Connell et al. 2014; De Martino et al. 2015) may shed light on the computational relevance of the cortical depth dependent organization of the auditory cortex (Moerel et al. 2018, 2019). The approach presented here can be followed for such future endeavors.

## Funding

This work was supported by the Dutch Research Council (NWO; grant 451-15-012 to M.M., and grant 864-13-012 to F.D.M.) and the European Research Council (ERC) under the European Union’s Horizon 2020 research and innovation programme (grant ERC-CoG-2020-101001270 to F.D.M).

## Competing interests

The authors declare no financial or non-financial competing interests.

## Author Contributions

M.M., F.D.M, and G.G. designed the research. M.M., F.D.M., and O.F.G. performed the research.

M.M. and A.L.C. analyzed the data. All authors contributed to writing the paper.

## Notes

### Competing Interest Statement

The authors have declared no competing interest.

