## Supplementary Material for "Selective attention reduces responses to relevant sounds in human auditory cortex"

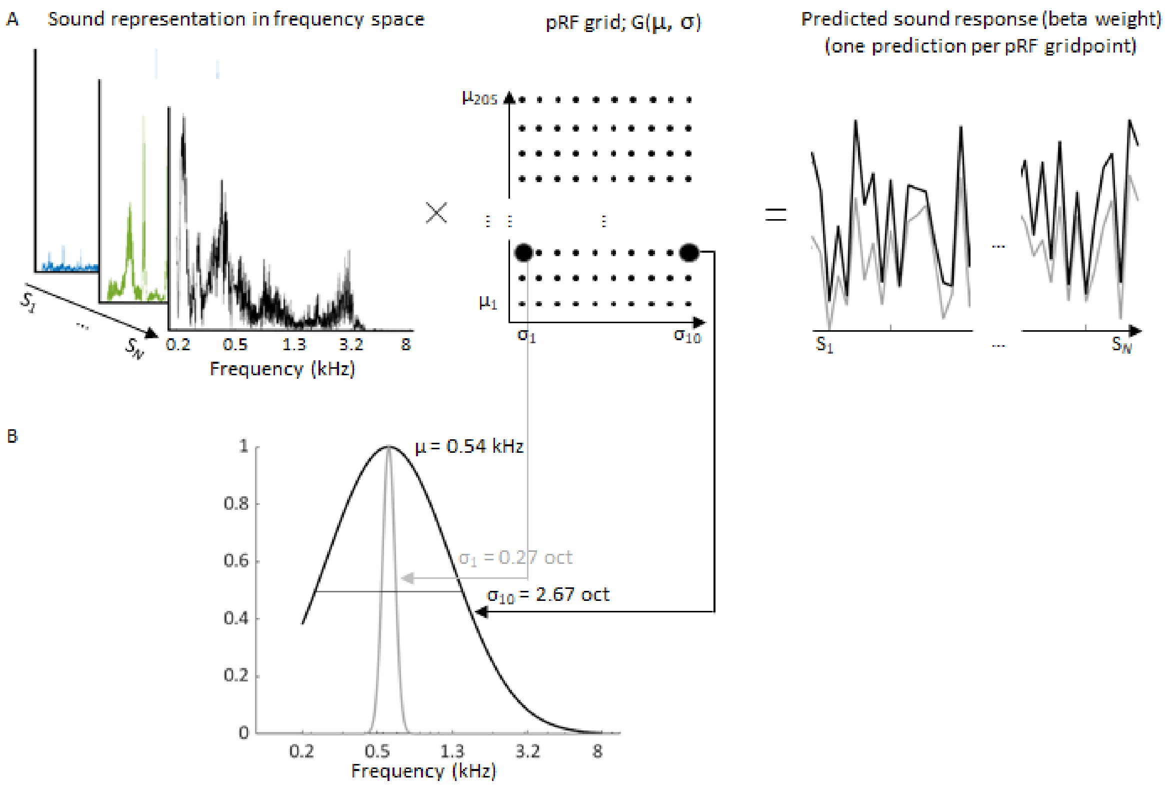


**Supplementary Figure S1. Schematic of the pRF estimation procedure.**

A) Sounds are represented according to their frequency content, with the frequency axis ranging from 180 to 8000 Hz in 2048 logarithmically-spaced bins. Responses to these sounds are predicted by multiplying each sound spectrum by a one dimensional Gaussian, defined by mean μ and size σ. The pRF grid defines the possible mu and sigma space (with μ ranging from 180 to 8000 Hz in 205 logarithmically-spaced bins, and σ ranging from 0.27 to 2.67 octaves in 10 logarithmically-spaced bins). B) Two example one-dimensional Gaussian pRFs, with the same frequency preference (μ = 0.54 kHz) but different selectivity (the most narrow tuning width of σ = 0.27 octaves is displayed by the gray line, and the broadest tuning width σ = 2.67 octaves is displayed by the black line). Tuning width is defined as the full-width-at-half-maximum (FWHM), indicated by the black dotted line. Note that while the pRF grid sampled the frequency space relatively coarsely (205 possible values of μ), the resulting pRF finely sampled the frequency space (2048 frequency bins).

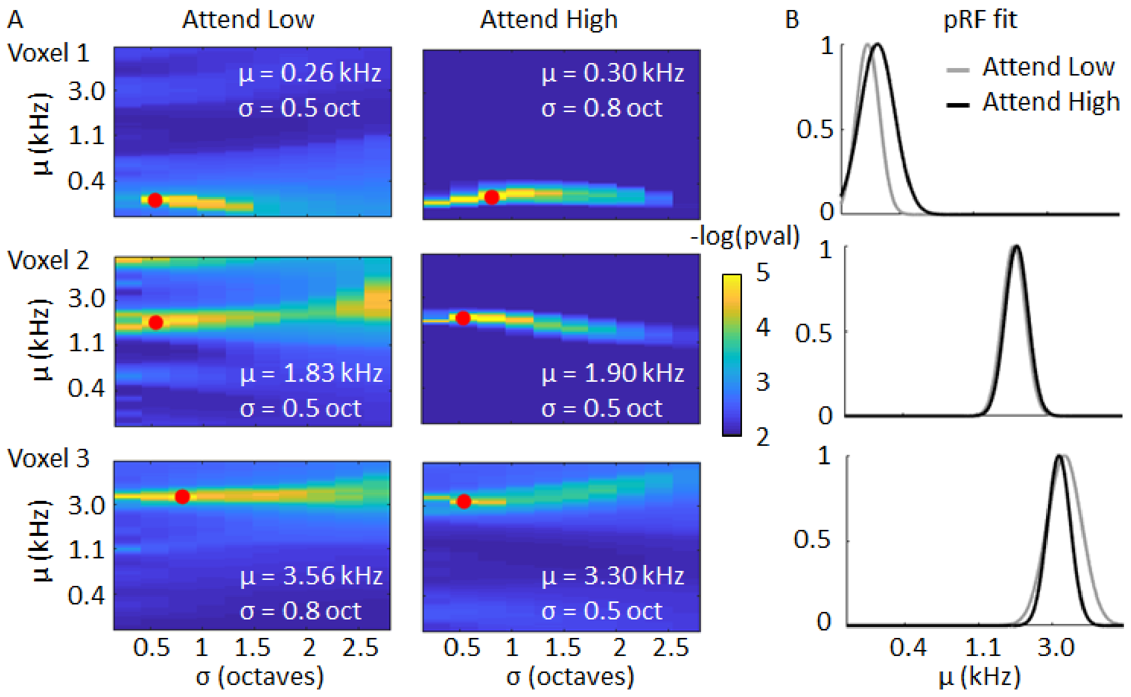


**Supplementary Figure S2. pRF fit.**

A) For three example voxels, the value of the cost function (-log(p value)) for all tested combinations of the pRF mean μ (BF in kHz) and size σ (TW in octaves) is shown. The red dots reflect the global minimum of the cost function, and correspondingly the BF and TW assigned to the voxel. B) The gray and black lines show the selected pRFs for the three example voxels per attentional condition. In these selected voxels, BF is stable across conditions while TW narrows with attention in those voxels whose BF is close to the attended one.

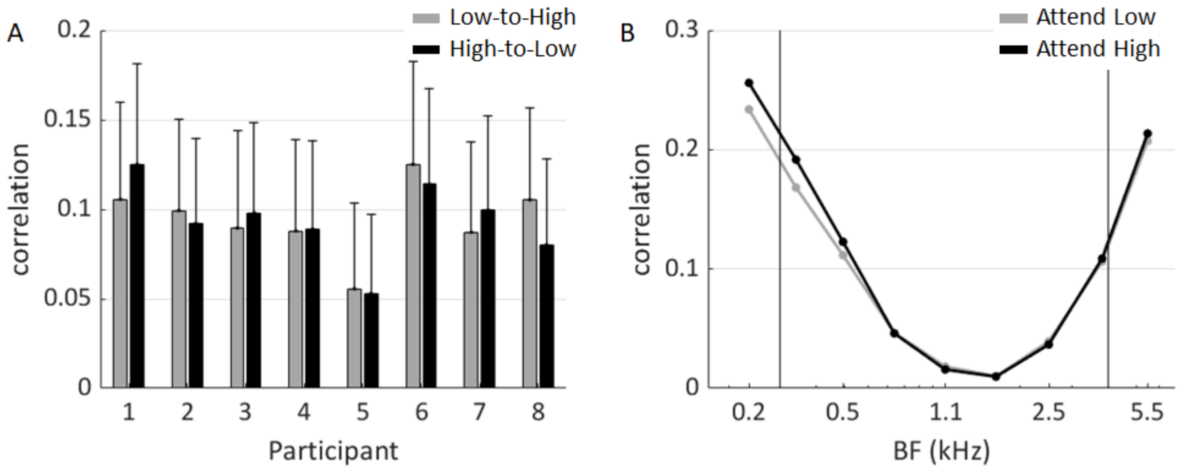


**Supplementary Figure S3. Prediction accuracy.**

Prediction accuracy as the Pearson correlation coefficient between predicted and observed responses to test sounds. All voxels with significant response to sounds and a prediction accuracy > 0.18 (corresponding to $p < 0.01$) in the “Baseline” dataset were included. A) Across-condition prediction accuracy (e.g., “Low-to-High” means predicting responses to test sounds in “Attend High” based on responses to training sounds in “Attend Low”), averaged across voxels and cross validations, in each participant. Gray and black bars show results from the “Low-to-High” and “High-to-Low” condition, respectively. Error bars represent the standard deviation across cross validations. B) Within-condition prediction accuracy as a function of the voxels’ best frequency (BF), averaged across cross validations and participants. Gray and black lines show results from the “Attend Low” and “Attend High” condition, respectively. The attended frequencies are indicated by black vertical lines.

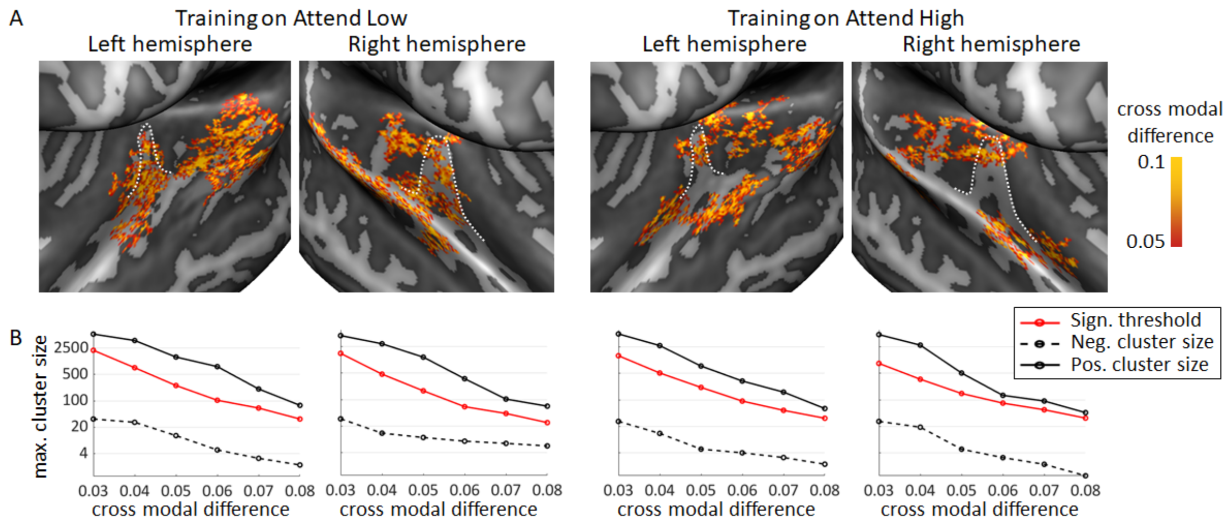


**Supplementary Figure S4. Cross modal prediction accuracy difference per attentional condition.**

A) Both when PRF fitting (i.e., model training) was based on data of the Attend Low condition (left) and when this was based on the Attend High condition (right), the cross modal difference (computed as [within-condition – across-condition prediction accuracy]) maps show positive values, indicating that prediction accuracy was higher when probed within- compared to across-attentional conditions. The white dashed line outlines HG. Maps are cluster size corrected at a FWER = 0.05. B) Across training conditions (left vs. right), hemispheres, and map thresholds ("cross modal difference"), the largest positive cluster size observed (solid black line) was greater than expected by chance (red line). The largest negative cluster size observed (dashed black line) did not exceed the cluster size expected by chance (red line).

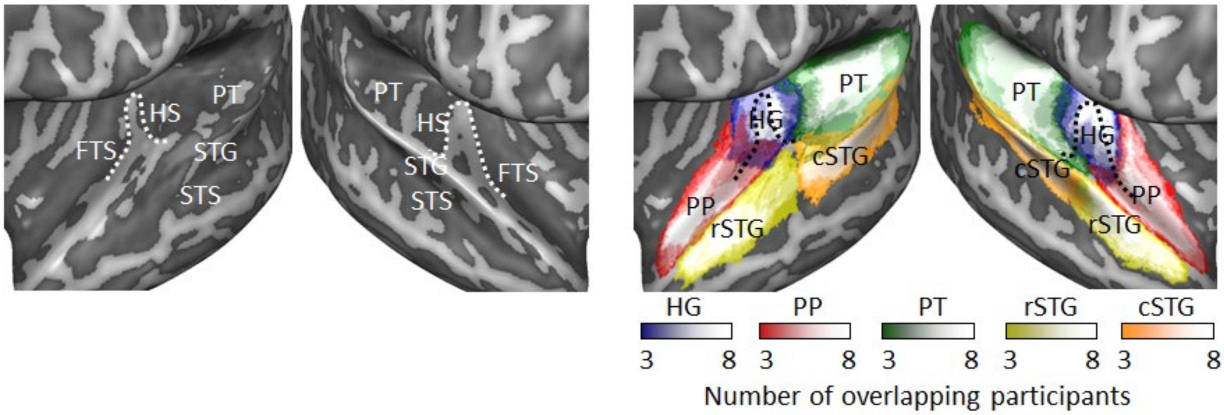


**Supplementary Figure S5. Regions of interest.**

(Left) Anatomy of superior temporal plane in the left and right hemisphere, visualized as the inflated group cortical surface mesh. The white dotted line outlines Heschl's gyrus (HG). FTS = first temporal sulcus; HS = Heschl's sulcus; PT = planum temporale; STG = superior temporal gyrus; STS = superior temporal sulcus. (Right) Across-participant overlap in regions of interest. Overlap from three to eight participants is shown ranging from dark to light hues. HG, planum polare (PP), PT, and rostral and caudal STG (rSTG and cSTG) are indicated in blue, red, green, yellow, and orange, respectively. The black dotted line outlines HG.

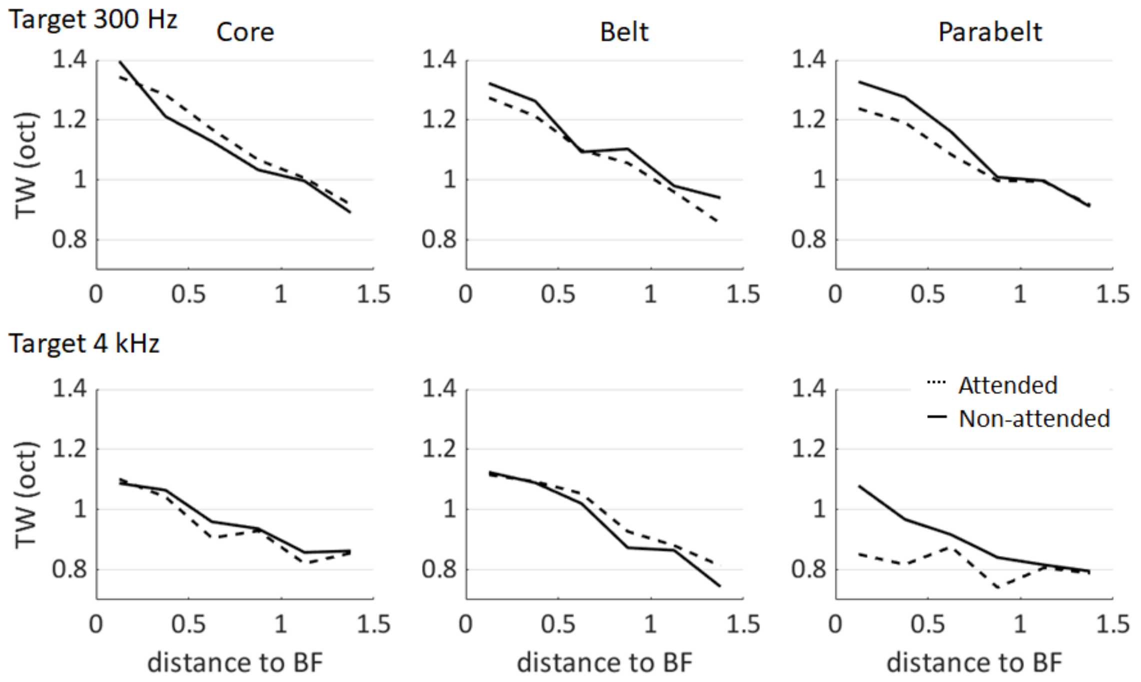


**Supplementary Figure S6. Effect of selective attention on tuning width per region and target frequency.**

Tuning width (TW) as a function of distance (in octaves) between BF and the attended target frequency, separate for when the target frequency was 300 Hz (top row) and 4 kHz (bottom row). For both target frequencies, TW in the parabelt region was more narrow when the BF was attended (dashed black line) compared to when it was not attended (solid black line). This effect was descriptively strongest in those voxels whose BF was closest to the attended sound frequency.
